# MicroRNA775 targets a *β-(1,3)-galactosyltransferase* to regulate growth and development in *Arabidopsis thaliana*

**DOI:** 10.1101/2021.01.28.428559

**Authors:** Parneeta Mishra, Akanksha Singh, Ashwani Kumar Verma, Rajneesh Singh, Sribash Roy

## Abstract

MicroRNAs are critical regulators of gene expression in plants and other organisms, and are involved in regulating plethora of developmental processes. Evolutionarily, miRNAs can be ancient and conserved across species or recently evolved and young, which are not conserved across diverse plant groups. miR775 is a non-conserved miRNA identified only in *Arabidopsis thaliana*. Here, we investigated the functional significance of miR775 in *A. thaliana* and observed that miR775 targets a probable *β-(1,3)-galactosyltransferase* gene at post transcriptional level. Phenotypic analysis of miR775 over-expression lines and the target mutant suggested miR775 regulates rosette size by elongating petiole length and increasing leaf area. Further, the expression of miR775 was found to be up-regulated in response to UV-B and hypoxia. Our results also suggest that miR775 regulated β-(1,3)-galactosyltransferase may involve in regulating the β-(1,3)-galactan content of arabinogalactans. Collectively, our findings establish a role of miR775 in regulating growth and development in *A. thaliana.*

**Highlights:**

- The role of an uncharacterized microRNA, miR775 has been explored
- miR775 targets a probable β-(1,3)-galactosyltransferase involved in complex carbohydrate biosynthesis
- miR775 regulates rosette size in *A. thaliana* and may play role under UV light and hypoxia

## 1. Introduction

MicroRNAs (miRNAs) belong to a class of small RNA of 20-24 nucleotides in length. They are processed from stem loop like structures of primary transcript, transcribed by RNA polymerase II to mature form by the activity of DCL1 protein. After processing miRNAs are loaded onto the ARGONAUTE 1 protein. The ARGONAUTE 1 loaded miRNA constitutes the RNA induced gene silencing complex (RISC) and carries out either post-transcriptional gene silencing or translational inhibition (Reinhart et al., 2002; Bartel, 2004; Vaucheret, 2008; Voinnet, 2009). In the past two decades, biological relevance of numerous miRNA in plants has been deciphered by observing the effect of miRNA over-expression or null expression, and also by portrayal of the target genes (Wang et al., 2005; Koyama et al., 2010; Marin et al., 2010; Zhao et al., 2015; Chung et al., 2016; Koyama et al., 2017). There exists plethora of information supporting the instrumental role of miRNA in plant stress response (Sunkar and Zhu, 2004; Lv et al., 2010; Zhou et al., 2010; Eckardt, 2012), growth and development (Wu and Poethig, 2006; Wu et al., 2006; Nag et al., 2009; Koyama et al., 2010; Marin et al., 2010; Rodriguez et al., 2010), signal transduction (Achard et al., 2004; Guo et al., 2005; Mallory et al., 2005), nutrient homeostasis (Chiou et al., 2006; Liang et al., 2010; Kawashima et al., 2011) and in regulating their own biogenesis (Xie et al., 2003; Vaucheret et al., 2004). However, majority of studies are focused on ancient miRNAs that are conserved across diverse plant species.

Despite the fact that miRNAs are well conserved between diverged plants, the vast repertoire of miRNA in plants is largely composed of non-conserved miRNAs. According to the reports, most of the non-conserved miRNAs are either restricted to a particular lineage or species. This gives an impression that a significant proportion of miRNA genes are young that have evolved much later in the evolutionary past. In disparity with the conserved miRNAs that largely target transcriptional regulators, the non-conserved miRNA are reported to target wide range of genes involved in varied plant processes (Rajagopalan et al., 2006; Fahlgren et al., 2007; Cuperus et al., 2011). For example, miR163 that is only reported in *Arabidopsis* has been identified to target members of a methyltransferase family, while miR158 which is confined only to the family Brassicaceae is known to target genes expressing PPR proteins (Chung et al., 2016; Ma et al., 2017). miR857 that is only encoded in *Arabidopsis* has been found to influence the growth of vascular tissue by targeting *Laccase 7* (Zhao et al., 2015). Besides targeting protein coding genes, some non-conserved 22 nt-miRNAs, for example miR173 targets non-coding RNA, triggering phasi RNA generation in *A. thaliana* and *A. lyrata* (Montgomery et al., 2008; Felippes and Weigel, 2009; Chen et al., 2010; Cuperus et al., 2010). Thus newly evolved miRNAs can have a considerable impact in managing the intricate processes in the plant body. Nevertheless, there exists discrepancy over the functional involvement of young miRNAs into the plant regulatory network. Studies suggest that it is not necessary that all the newly identified miRNAs have gained a function. Some may be in the evolutionary transient phase and disappear during the course of evolution (Rajagopalan et al., 2006; Fahlgren et al., 2007; Cuperus et al., 2011). Thus the function of newly evolved species specific miRNAs remains largely unexplored.

miR775 is one such young uncharacterized miRNA reported so far only from *A. thaliana* (Lu et al., 2006; Rajagopalan et al., 2006). The precursor of this miRNA maps on chromosome 1 and is 123 nucleotides (nts) long that is processed into its 20-nts long mature form. In our previous study, we observed that the expression of miR775 was higher in the high altitude populations of *A. thaliana* (Chit and Mun) as compared to the lower altitude population (Deh) under both native and control conditions (Tripathi et al., 2019) (Supplementary Table S4). Thus, we speculated that miR775 might have a significant role in *A. thaliana*. We show that miR775 targets a probable β-(1,3)-galactosyltransferase belonging to glycosyltransferase 31 (GT31) family of carbohydrate active enzymes that might be involved in regulating arabinogalactan and/or complex carbohydrate biosynthesis, an important component of plant cell wall. We also show increased expression of miR775 under UV light and hypoxic conditions, and thus might help plants under such abiotic stress conditions, mostly prevalent in high elevated areas. However, during our reporting time of this work Zhang et al. (2021) also reported role of miR775 in *A. thaliana.*

## 2. Material and methods

### 2.1. Plant material

*A. thaliana* ecotype, Columbia-0 (Col-0) was used for all the experimental purposes. Seeds of T-DNA insertion line for target knockout mutant (SALK_151601) and miRNA mutant (SALK_043658) were obtained from Arabidopsis Biological Resource Center (ABRC, USA). *A. thaliana* plants were grown at 22°C under controlled conditions with 140μmol/m^2^/sec light intensity and 16-hours-light/8-hours-dark photoperiod cycle. Seeds were stratified for 4 days at 4°C in dark before being transferred to the controlled culture room. The SALK lines were first confirmed for homozygosity using the primer combination of LBb1.3+LP+RP as discribed in GABI (http://signal.salk.edu/tdnaprimers.2.html), and then using qRT-PCR for functional confirmation. Preliminary screening indicated that the T-DNA inserted line SALK_043658 used as probable miR775 mutant was not an effective mutant and hence was not used in further studies.

### 2.2. RNA isolation and gene expression analyses by qRT-PCR

Total RNA was isolated using *mir*Vana™ miRNA/ total RNA isolation kit (Ambion, USA) as per manufacturer’s protocol. RNA integrity was checked by resolving 2μl of total RNA on 1% agarose gel. The RNA was subjected to DNase (Turbo DNaseI) treatment and cDNAs was synthesized from one microgram (μg) of total RNA with miR775 stem loop primer using Super-Script III reverse transcriptase (Invitrogen, USA). To check the expression level of miR775, cDNAs were amplified using miR775 specific forward primer and stem loop specific universal reverse primer. For mRNA expression analysis, qRT-PCR was performed using cDNA synthesized from one μg of total RNA with oligo-dT using First Strand cDNA synthesis kit (Invitrogen, USA) according to manufacturer’s instructions and PCR amplification was done using gene specific primers.

qRT-PCR was performed using ABI SYBR Green (Applied Biosystem, USA) on ABI7300 real-time PCR system (Applied Biosystem, USA) with the following PCR conditions: denaturation at 95°C for 10 mins, followed by 40 cycles of denaturation at 95°C for 20sec, annealing and extension together at 60°C for 60sec. The expression level was determined following 2^−ΔΔCt^ method (Schmittgen and Livak, 2008). The plant *actin* gene and *5S rRNA* gene were taken as internal control for normalization of mRNA and miRNA expression respectively. The primer sequences used are listed in Supplementary Table 1.

### 2.3. *In-silico* target prediction and its validation

The targets of miR775 were predicted using psRNATarget prediction tool (http://plantgrn.noble.org/psRNATarget/) at default parameters. Based on target prediction, the putative target of miR775 was validated by modified 5′RLM-RACE using First Choice RLM-RACE Kit (Ambion,USA) following manufacturer's instructions with minor modifications. Briefly, five μg of total RNA isolated from rosette tissues was subjected to adapter ligation at 37°C for 1 hour, followed by reverse transcription at 42°C for 1 hour. The prepared cDNA was used as template for outer RACE PCR using outer_adapter primer (provided by the manufacturer) and outer_GALT primer (target gene specific primer). Nested PCR was further carried out with inner_adapter primer (provided by the manufacturer) and inner_GALT primer (target gene specific primer). The PCR product was run on 2% low melting agarose gel and the PCR fragment was excised, eluted and cloned in pGEM-T easy vector (Promega, USA). Further 12 individual clones were sequenced for analysis and confirmation of the target cleavage site.

### 2.4. Preparation of different constructs and generation of transgenic lines

To over-express miR775, region flanking miR775 precursor (555bp) was amplified from Col-0 genomic DNA using MIR775_PRI_forward and MIR775_PRI_reverse primers (Supplementary Table 1). The amplicon was sub cloned in pGEM-T easy vector (Promega, USA) for sequencing, and then transferred into plant expression vector pBI121 between the CAMV35S promoter and Nos terminator following digestion with *Xba*I and *Sa*cI. For promoter study, the 560 bp region upstream of the transcription start site was amplified using Q5 high fidelity DNA polymerase (NEB, England) and ligated into *EcoR*V digested SK+ cloning vector. The transcription start site (TSS) for *MIR775* primary transcript was identified using AtmiRNET resource available online (http://atmirnet.itps.ncku.edu.tw/promoter.php.). Post sequence confirmation, the promoter region was excised from the cloning vector by sequential digestion with *Sma*I and *Xba*I, and cloned into binary vector pBI101 just upstream of the *GUS* reporter gene. The binary plasmid harbouring the constructs were individually transformed in *Agrobacterium* cells, GV3101 and plant transformation was done by floral dip method following Clough and Bent (1998). Putative transformants were first screened on MS agar plates containing kanamycin (50ug/ml) and further by PCR.

### 2.5. Phylogenetic analysis of the target gene of miR775

Phylogenetic analysis of the target gene of miR775 was performed using online tool Phylogeny.fr. (http://www.phylogeny.fr/simple_phylogeny.cgi). Phylogeny was generated following ‘one click’ mode that functions on a pre-set pipeline consisting of MUSCLE for sequence alignment, G-block for alignment curation, PhyML and Tree Dyn for tree formation (Dereeper et al., 2008). Protein sequence of all the members of GT31 family in *A. thaliana* were retrieved from TAIR database.

### 2.6. Analysis of *MIR775* promoter

*MIR775* promoter was analyzed by histochemical GUS staining in different tissues of *MIR775*PRO*:GUS* transgenic lines following standard method (Jefferson, 1989). Briefly, tissues were dipped in a standard solution containing 100 mM Na-phosphate buffer (pH 7.2), 10 mM EDTA, 2 mM potassium ferricyanide, 2 mM potassium ferrocyanide and 5-bromo-4-chloro-3-indolyl-β-D-glucuronide at a concentration of 1mg/ml and incubated for 16 hours at 37°C. Stained samples were incubated in 70% ethanol for chlorophyll removal, and images were taken on Leica EZ4E stereo microscope.

The *cis* regulatory motifs present in the promoter sequence of *MIR775* were identified using online tool Plant CARE (http://bioinformatics.psb.ugent.be/webtools/plantcare/html/).

### 2.7. High light and UV-B treatment

To examine the effect of high light intensity on expression of miR775, 21 days old plants were exposed to 600μmole/m^2^/sec or 1000μmole/m^2^/sec of light intensity for an hour. Leaf sample was harvested and frozen in liquid nitrogen for storage till RNA isolation. The effect of UV light on expression of miR775 was examined by exposing 21 days old plants to UV-B light (TL20W/01RS, Philips) in a close chamber maintaining a distance of 30 cm from the light source for an hour. Post treatment leaf tissue was collected and frozen in liquid nitrogen for storage till RNA isolation. Both high light and UV-B treatment experiments were conducted in triplicates.

### 2.8. Hypoxia treatment/Mitochondrial inhibitor treatment

Hypoxia treatment was performed by treating plants with mitochondrial inhibitors following Bond et al. (2009) with minor modifications. Briefly, the plants were sprayed with 50 μM antimycin A (AA, Sigma) and 5 mM salicylhydroxamic acid (SHAM, Sigma) ensuring uniform spraying on all the plants. The experiment was started one hour before the onset of day light, and spraying was repeated after every hour. Sampling for RNA isolation was done three hours after treatment by pooling leaf tissue from 5 individual plants. For estimation of hydrogen peroxide, samples were collected six hours after the treatment. Hydrogen peroxide was detected by 3, 3’-diaminobenzidine (DAB) staining as described by Daudi and O’Brien (2012) and quantified following Alexieva et al. (2001). Each experiment was conducted in triplicate.

### 2.9. Morphological traits analyses

Morphological variations amongst the wild type (WT) Col-0, miRNA over-expression lines (*MIR775*OXP) and target mutant (SALK_151601) were recorded. Plants were grown in pots containing soilrite under standard growth conditions (as described above). Both vegetative and reproductive traits including rosette diameter (major and minor), number of leaves, leaf length and width, petiole length, bolting time, plant height, total number of siliques, silique length, and number of seeds per silique were recorded from nine to twelve individual plants of each WT, *MIR775*OXP and SALK_151601. Data were plotted as means ± standard errors (SE). Statistical analysis was done using one way ANOVA followed by post hoc test to check the significance between the groups using R (Team, 2017).

### 2.10. β–Galactosyl Yariv staining for detection of arabinogalactans

β–Galactosyl Yariv (β–Gal-Yariv) staining in stem sections, and pollen grain was done following Popper (2011). Briefly, the stem sections were incubated in 50μl of β–Gal-Yariv reagent at a concentration of 2mg/ml in 0.15M NaCl with continuous shaking for an hour. Post incubation washing was done with 0.15M NaCl and samples were mounted in the same for observation under Leica DM2500 microscope.

### 2.11. Calcofluor white staining for detection of seed coat phenotype

Calcofluor white staining was performed following Willats et al. (2001). Briefly, seeds were imbibed overnight in water with continuous shaking at room temperature. Imbibed seeds were treated with 1ml calcofluor white solution for 10 mins, followed by repeated washing with water. Imaging was done using confocal microscope (Zeiss LSM 510) with excitations at 405 nm.

## 4. Results

### 4.1. miR775 is ubiquitously expressed in *A. thaliana*

We examined the expression pattern of miR775 in different tissues of *A. thaliana* using qRT-PCR. Expression of miR775 was detected in all the examined tissues including seedling, root, rosette leaf, cauline leaf, stem, inflorescence and siliques. However, expression of miR775 was relatively higher in stem, siliques and inflorescence as compared to other tissues, while in roots its expression was the least (Fig. 1A). To further ascertain the expression pattern in different tissues, we cloned the putative promoter of *MIR775* (560bp upstream of TSS) and fused it with *GUS* reporter gene. Transgenic lines expressing *GUS* driven by *MIR775* promoter (*MIR775*PRO-*GUS* lines) were generated and analyzed by GUS staining. The GUS expression was observed in various tissues at different developmental stages (Fig.1B). GUS expression was largely confined to hypocotyls, petiole, root tips and veins of young leaves, adult rosette leaf and cauline leaf. Expression of GUS was limited to anthers and stigma of flower tissues as observed with higher expression of miR775 in inflorescence. Data of qRT-PCR and GUS staining indicates miR775 expresses ubiquitously in *A. thaliana.*

**Fig. 1:**
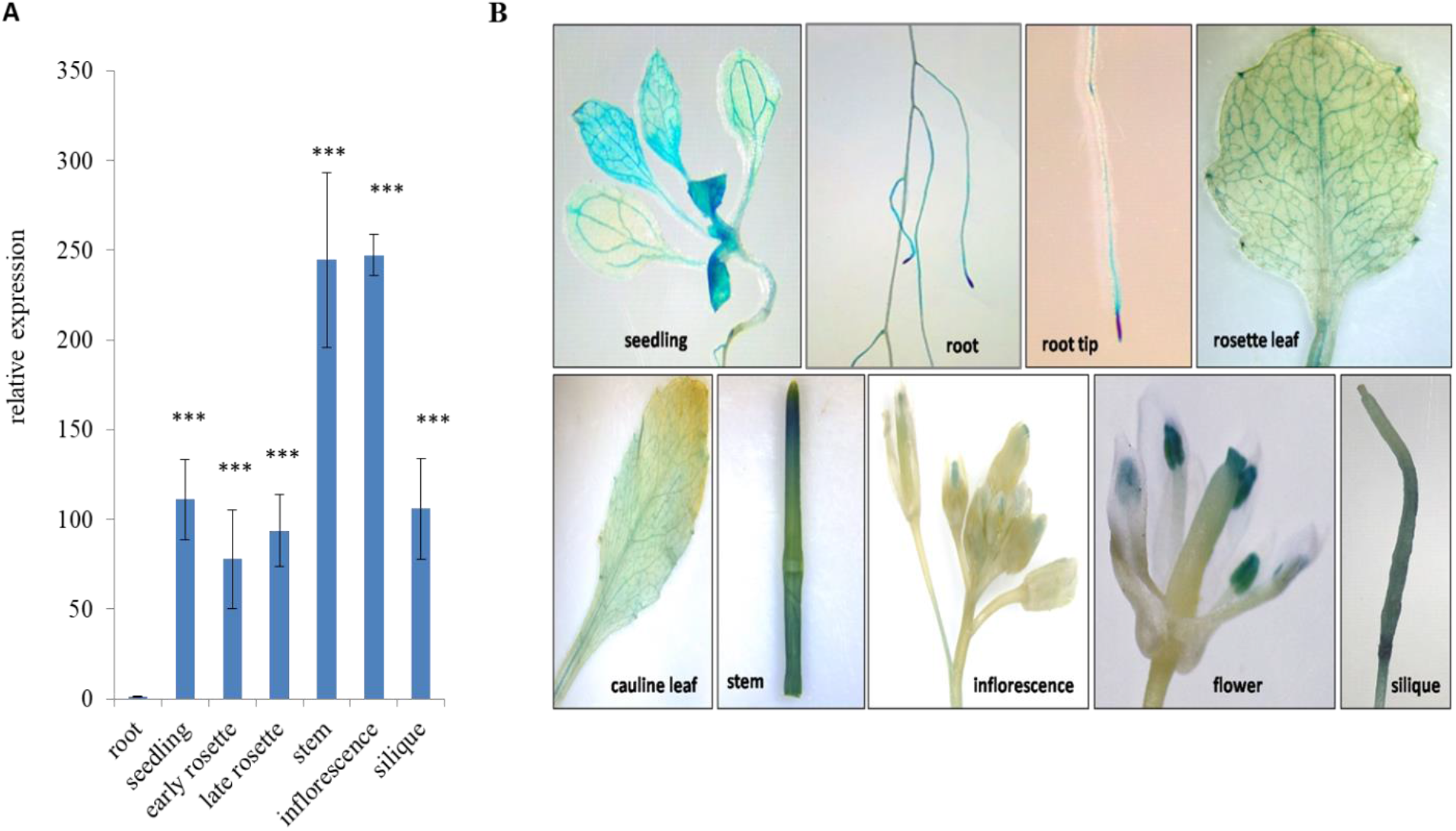
Characterization of miR775 in *A. thaliana* (A) Relative expression of miR775 in different tissues of WT Col-0 as determined by qRT-PCR. Seedling tissue was taken from 1 week old plants, early and late rosette were taken from 15 and 30 days old plants respectively, while root, stem, inflorescence and siliques were collected from 45 days old plants. Data represents mean of three replicates ±SE, *** indicate values which were significantly different at P value <0.001 (Student’s t-test). (B) Histochemical GUS staining in different tissues of *MIR775*PRO*-GUS* transgenic lines. Seedling and roots tissue were taken from 15 days old plants grown on MS media, rosette leaf was taken from 30 days old plants while cauline leaf, stem, inflorescence, flower and silique were collected from 45 days old plants grown in soilrite.

### 4.2. miR775 targets a probable *β-(1,3)-galactosyltransferase* gene

*In-silico* target prediction analysis suggested 12 putative targets of miR775 including both cleavage and translational inhibition mode of operation (Supplementary Table 2). Amongst these, the gene encoding galactosyltransferase (GALT) protein (AT1G53290.1), was likely to be the most accurate target of miR775 according to the best target identification criteria (Dai and Zhao, 2011). Earlier report also predicted it to be the target of miR775 (Fahlgren et al. 2007). Therefore, we attempted to experimentally validate this target by modified 5'RLM-RACE. Target validation using modified 5'RLM-RACE identified the gene encoding GALT protein (AT1G53290) as the only target of miR775. The maximum numbers of detected transcripts (nine of 12 clones) were cleaved at the expected position (525 nts from 5’ end the mRNA) i.e. between the 10^th^ and 11^th^ nucleotide (nt) from 5’ end of miRNA-target pair (Fig. 2A), confirming cleavage of *GALT* transcript by miR775. We also attempted to validate *DCL1*, another predicted target of miR775 but we could not amplify the expected fragment size by PCR during modified 5’RLM-RACE.

**Fig. 2:**
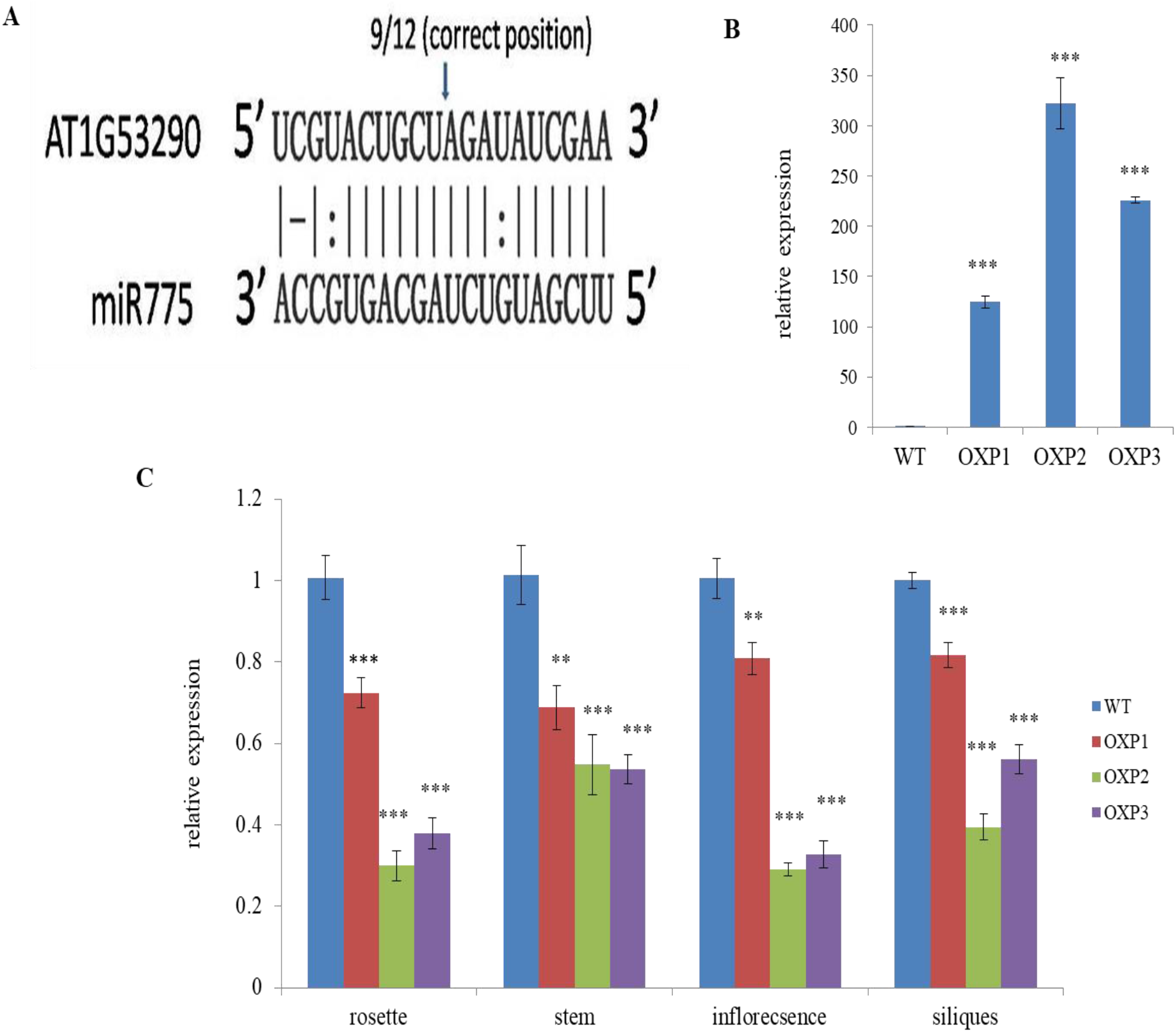
Targeting of AT1G53290 by miR775 (A) Cleavage of AT1G53290 mRNA by miR775 as detected by modified 5’RLM-RACE is shown. The fraction of clones with cleavage at expected 5’end to the total number of clones sequenced is indicated above the target site. (B) qRT-PCR analyses of miR775 in rosette tissue of homozygous *MIR775*OXP lines and WT (C) qRT-PCR analyse of *galactosyltransferase* target gene (AT1G53290) in rosette, stem, inflorescence and siliques of homozygous *MIR775*OXP lines and WT. Data is a mean of three replicates ±SE. **, *** indicate values which were significantly different from WT at P value <0.01, <0.001 respectively (Student’s t-test).

In order to gain further insight into the biological and functional significance of miR775, transgenic lines over-expressing miR775 (*MIR775*OXP) under the constitutively expressed CaMV35S promoter were developed and analyzed. qRT-PCR analysis of three homozygous *MIR775*OXP lines showed more than 100 fold higher expression of miR775 as compared to the WT (Fig. 2B). Subsequently, we determined the expression of the target *GALT* in different tissues of the *MIR775*OXP lines. qRT-PCR analyses suggested expression of target *GALT* was down-regulated in all the *MIR775*OXP lines as compared to the WT, in all the examined tissues (Fig. 2C). As expected, the OXP line having less expression of miR775 exhibited minimum target down-regulation. Further, to rule out that *DCL1* was not a true target of miR775, we analysed its expression and did not observe significant down-regulation in any of the *MIR775*OXP line with respect to WT (Supplementary Fig. 1). We also checked the expression pattern of the target *GALT* in different tissues in WT. Expression of target *GALT* though was found to be ubiquitous, but its expression was comparatively higher in stem, inflorescence and siliques than other tissues as observed with miR775 (Supplementary Fig. 2). The high level expression of both miRNA and the target gene in the same tissue have been reported earlier in other miRNAs as well (Zhang and Li, 2013; Sharma et al., 2016).

The target GALT is reported to be a GT31 family member of carbohydrate-active-enzymes (http://www.cazy.org) (Strasser et al., 2007; Qu et al., 2008). Moreover, recently a few galactosytransferases have been characterized. We re-examined the phylogenetic relationship of the miR775 target GALT with other similar protein sequences. A simple protein BLAST with target GALT at NCBI identified a number of proteins with sequence similarity ranging from 72 to 100 percent and query coverage of 98 to 100 percent. Phylogenetic analysis of these proteins (98) revealed that the target GALT of miR775 was closely related to β-(1,3)-galactosyltransferases of *Camelina sativa* and *Arabidopsis lyrata* (Fig. 3A). Phylogenetic analyses with 33 *A. thaliana* GT31 members placed it near the clade containing two β-(1,3)-galactosyltransferases (AT1G87100 and AT1G33430) and one β-(1,6)-galactosyltransferase (AT1G32930), those are reported to be involved in arabinogalactan biosynthesis (Qu et al., 2008; Geshi et al., 2013; Suzuki et al., 2017) (Fig. 3B), although other members of the clade are yet to be characterized.

**Fig. 3:**
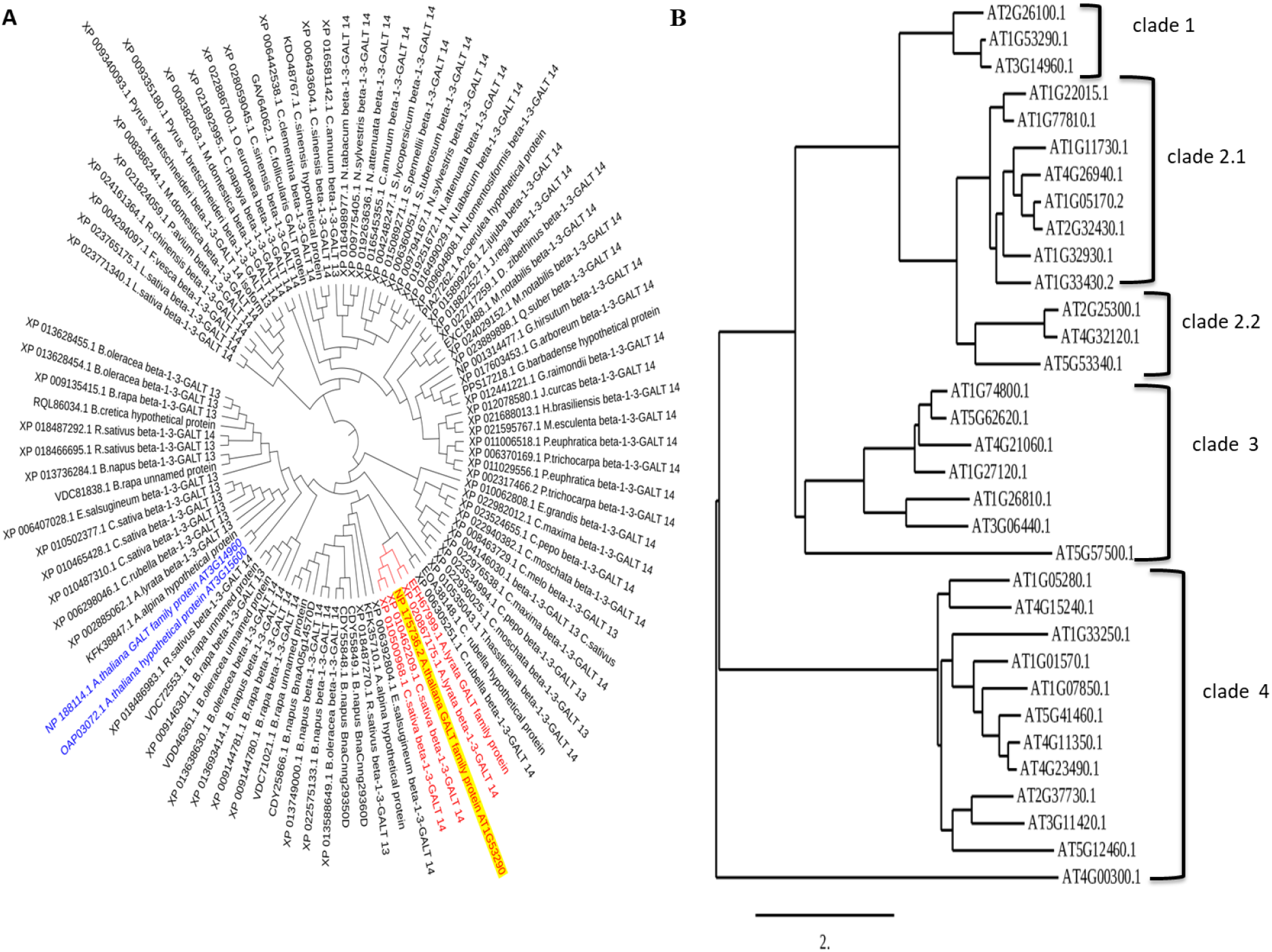
Characterization of miR775 target gene (A) Phylogenetic analysis of AT1G53290 on the basis of sequence similarity with other plant species (B) Phylogenetic analysis of AT1G53290 with *A. thaliana* members of GT31 family. All the members of clade 4 and AT5G57500.1 of clade 3 lack the GALT domain (Qu et al., 2008). All other members of clade 3 are hydroxyproline GALTs (Basu et al., 2015a; Basu et al., 2015b), except AT1G26810.1 which is a β-(1,3)-GALT involved in N-glycan biosynthesis (Strasser et al., 2007). All members of sub-clade 2.2 are hydroxyproline GALTs (Ogawa‐Ohnishi and Matsubayashi, 2015). Members of sub clade 2.1, AT1G87100 and AT1G33430 are β-(1,3)-GALTs while AT1G32930 is β-(1,6)-GALT (Qu et al., 2008; Geshi et al., 2013; Suzuki et al., 2017). Other members of sub clade 2.1 and all members of clade 1 are awaiting functional validation.

### 4.3. miR775 is UV-B and hypoxia responsive

Since the expression of miR775 was higher in high altitude populations, we hypothesized that the expression of miR775 might be governed by environmental factors like high light, UV light etc. prevalent in those areas. *In-silico* analysis of *MIR775* promoter revealed the presence of several regulatory motifs including light, hypoxia etc. (Supplementary Table 3, Fig. 4A). Three types of light responsive motifs were detected, namely the ACE (CTAACGTATT) element, Box 4 (ATTAAT) and I box (cCATATCCAAT). The ACE element is reported to facilitate HY5 mediated transcriptional regulation of genes in response to light and UV-B radiation (Logemann and Hahlbrock, 2002; Safrany et al., 2008; Stracke et al., 2010; Huang et al., 2012; Gangappa and Botto, 2016; Qian et al., 2019). The I box motif is a part of light responsive element and is reported to be present in the upstream regulatory sequences of light regulated genes (Giuliano et al., 1988; Houston et al., 2013; Chao et al., 2014), and the Box 4 motif that is conserved across plants is involved in transcriptional regulation of genes associated with light stress (Chao et al., 2014). In *MIR775* promoter sequence, the Box-4 motifs were present at 238nts and 253nts upstream of the TSS, while other two light responsive motifs were present singly. However, when we exposed the WT plants to high light conditions (600μmol/m^2^/sec and 1000 μmol/m^2^/sec) for one hour, we could not detect any significant difference in expression level of miR775 (Fig. 4B). Further, considering the prevalence of UV-B in high altitude area and some of the light responsive elements are UV-responsive (like the ACE present in promoter of *MIR775*), expression of miR775 was analysed from UV-B treated plants. Interestingly, a significant increase in the expression of miR775 was observed in post UV-B treated plants (Fig. 4C).

**Fig 4:**
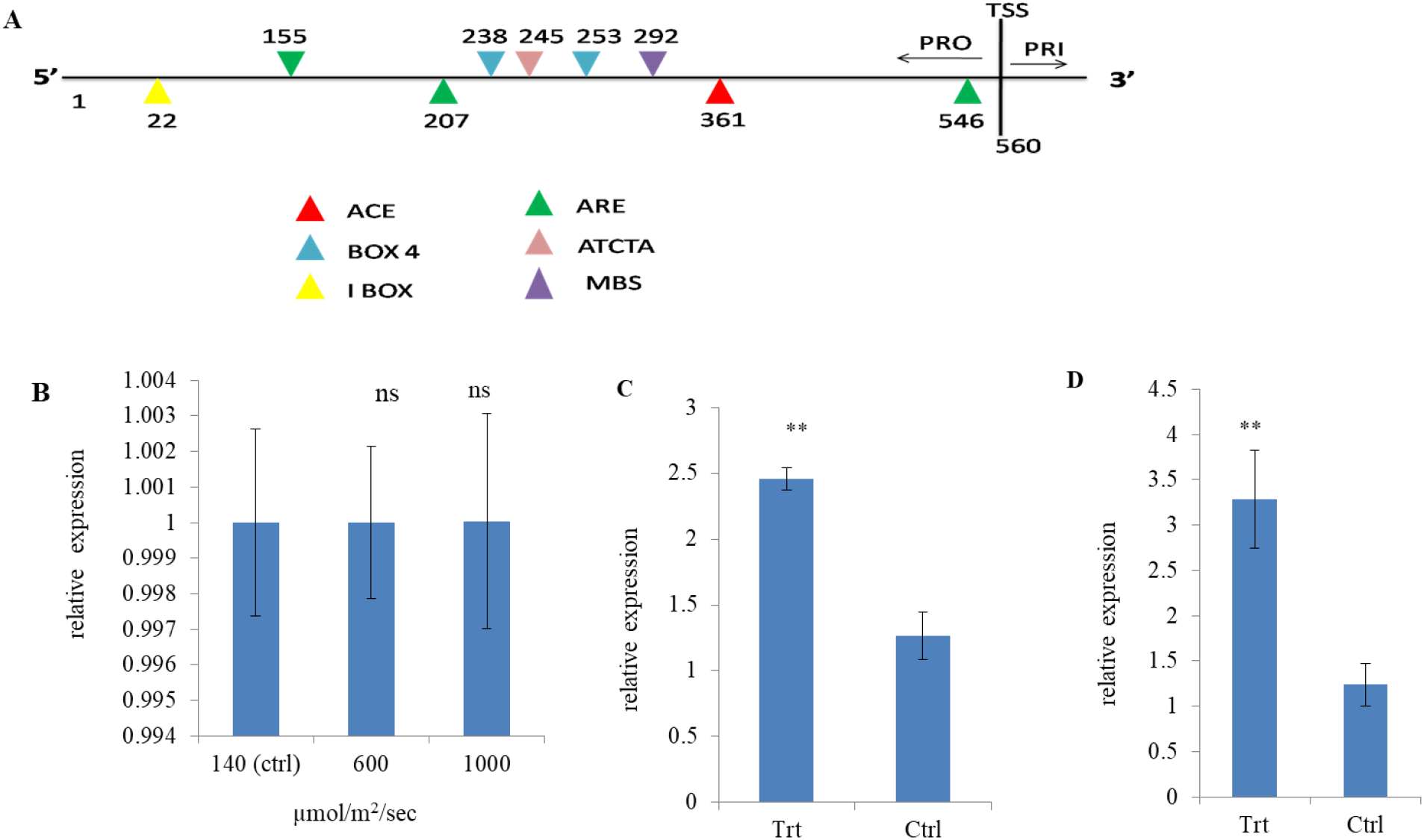
Expression of miR775 in response to environmental variables (A) Schematic representation of *MIR775* promoter showing regulatory motifs responsive to environmental factors. Real time expression analysis of miR775 (B) after 1 hour of high light treatment (C) after 1 hour of UV-B treatment (D) after 3 hours of treatment with mitochondria inhibitors, Antimycin A and Salicylhydroxamic acid. Data is a mean of three replicates ±SE. ** indicate values which were significantly different from control with P value <0.01 (Student’s t-test), ns indicates non-significant variation with respect to control.

Besides light responsive elements, there are three ARE and one ATCTA elements in the promoter region of *MIR775*. The ARE and ATCTA elements are reported to be present in the promoter of *Alchohol Dehydrogenase1,* a hypoxia responsive gene (Walker et al., 1987; Dolferus et al., 1994; Hinz et al., 2010). The presence of three ARE motifs, with one in the immediate vicinity of the TSS suggested a strong influence of this motif in governing *MIR775* promoter activity. We tested the expression of miR775 under hypoxic condition by treating the plants with mitochondrial respiratory inhibitors. The data suggested treatment with mitochondrial inhibitors led to an increase in miR775 level (Fig. 4D). Further, we determined the level of hydrogen peroxide under hypoxic conditions in WT, *MIR775*OXP and target mutant. After six hours of treatment with mitochondrial inhibitors, hydrogen peroxide accumulation was more in the target mutant and *MIR775*OXP as compared to the WT (Supplementary Fig. 4). This suggests that reduced level of the target gene in both target mutant and *MIR775*OXP promotes hydrogen peroxide accumulation in response to hypoxia.

### 4.4. miR775 affects rosette size

To determine the role of miR775 in plant growth and development, WT, *MIR775*OXP and target mutant plants were phenotypically characterized. The rosette area was significantly higher in *MIR775*OXP and the target mutant as compared to WT, 20 days post-germination (Fig. 5A, 5B, Supplementary Table 4). This was mainly due to expanded leaves with elongated petiole in *MIR775*OXP and the target mutant as compared to the WT (Fig. 5C, 5D). The rosette area of the target mutant was higher than the *MIR775*OXP. Interestingly, the number of rosette leaves was more in target mutant than WT, though there was no significant difference in rosette leaf number between *MIR775*OXP and WT (Fig. 5D). There were no significant variations in bolting time, plant height or total number of siliques in the target mutant and *MIR775*OXP as compared to WT. Although the target mutant had longer silique size and more number of seeds per silique than WT, but no difference was observed between *MIR775*OXP and WT (Fig. 5E). These results indicate prominent effect of miR775 in influencing rosette size, but relative miR775/target stoichiometry may not favour its tight regulation of reproductive traits.

**Fig. 5:**
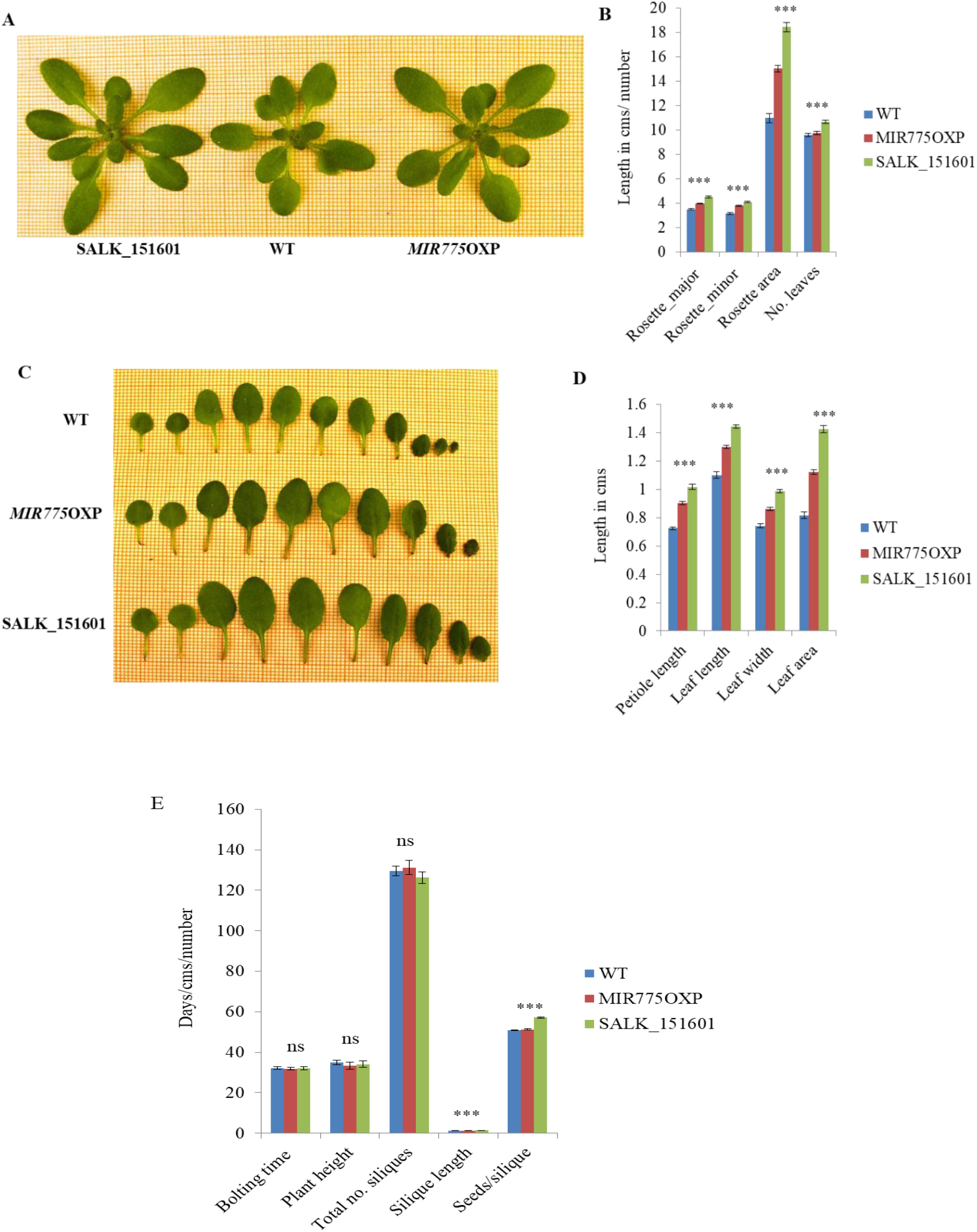
Analyses of morphological traits **(A)** image showing variation in rosette size **(B)** graphical representation of rosette parameters showing mean of WT, *MIR775*OXP, target mutant (SALK_151601). Error bars represent ±SE with n=12 **(C)** image showing variation in leaf dimensions **(D)** graphical representation of leaf parameters showing mean WT, *MIR775*OXP, target mutant (SALK_151601). Error bars represent ±SE with n=12. **(E)** Graphical representation of post vegetative traits showing mean of WT, *MIR775*OXP, target mutant (SALK_151601). Error bars represent ±SE with n=9. *** indicates significant variations amongst the groups with p value <0.001 according to One Way ANOVA, ns indicates non-significant variation.

### 4.5. miR775 targeted β-(1,3)-galactosyltransferase may be involved in complex carbohydrate biosynthesis

The β-(1,3)-galactosyltransferases are reported to be involved in the formation β-(1,3)-galactan content of arabinogalactan proteins (AGPs) and rhamnogalacturonan I (RG-I), a pectin glycan (Qu et al. 2008; Showalter and Basu 2016; Suzuki et al. 2017). Further, using STRING tool (https://string-db.org/) (Szklarczyk et al., 2015; Szklarczyk et al., 2019), the target β-(1,3)-galactosyltransferase was predicted to be functionally associate with proteins involved in protein glycosylation, complex carbohydrate biosynthesis and cell wall biogenesis (Supplementary Fig. 5, Supplementary Table 5). Thus it was speculated that the target GALT may be involved in regulation of the β-(1,3)-galactan content of arabinogalactans. The qualitative test of arabinogalactans using β-Gal-Yariv staining showed there was marked difference in staining intensity in stem cross sections of the WT and the null mutant of the target gene, but not between the *MIR775*OXP and WT (Fig. 6A, Supplementary Fig. 6A). However, there was no difference in β-Gal-Yariv staining intensity in the pollen grains of both target mutant and *MIR775*OXP as compared to WT (Supplementary Fig. 6B). Galactosyltransferases are reported to cause variation in seed coat phenotype by altering the properties of the seed coat mucilage (Basu et al., 2015a; Basu et al., 2015b). Therefore, the seed coat phenotype was analysed by staining with calcofluor white and visualized under confocal microscope. As evident the target mutant exhibited reduced staining as compared to WT and *MIR775*OXP (Fig. 6B). As observed with β-Gal-Yariv staining, there was no observable variation between *MIR775*OXP and WT on calcoflour white staining. Altogether, these results suggest that the target gene of miR775 is possibly involved in governing the biosynthesis of complex carbohydrates.

**Fig. 6:**
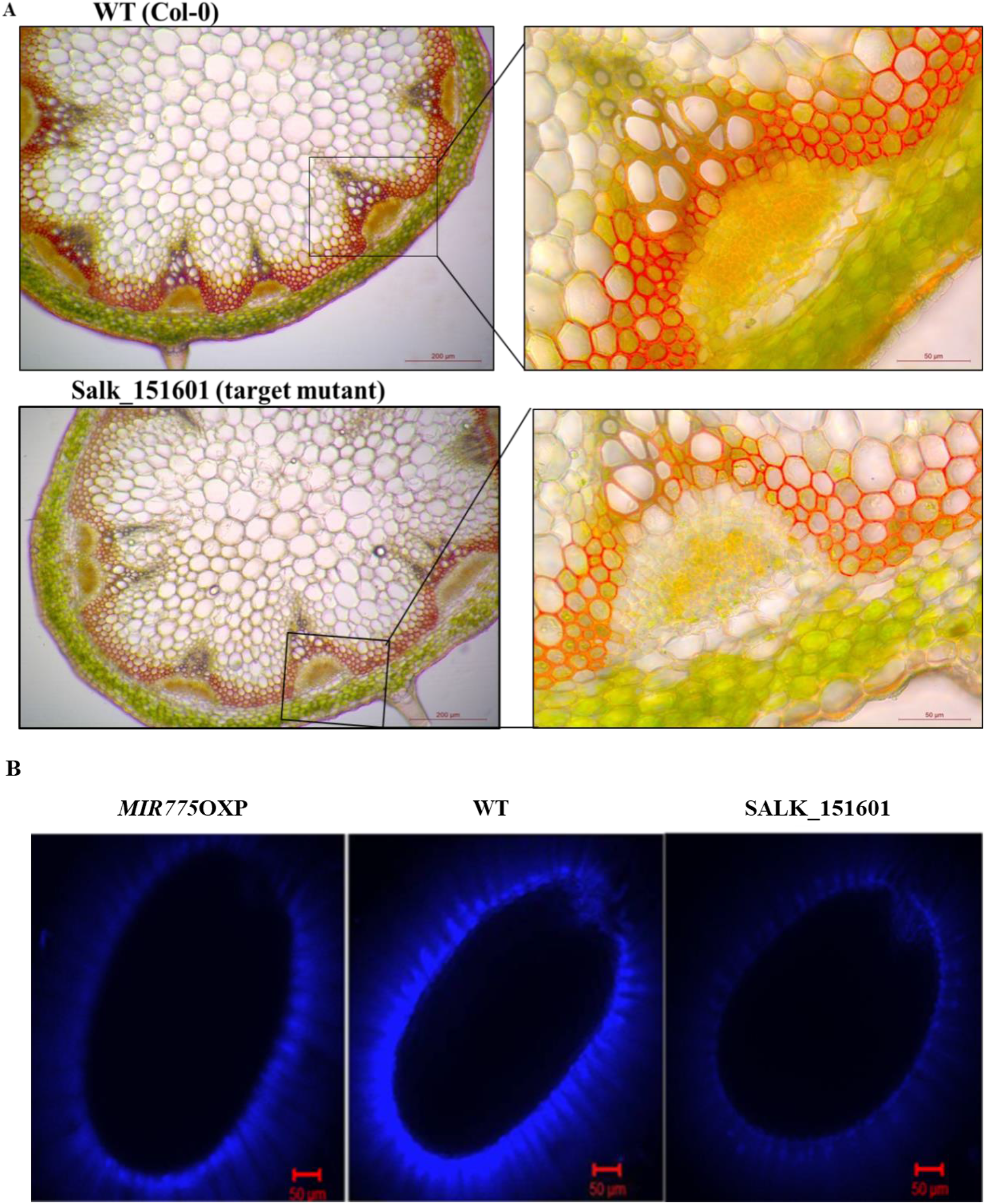
Involvement of miR775 target gene in complex carbohydrate biosynthesis (A) Staining with β-Gal Yariv reagent in stem sections of WT and target mutant (SALK_151601) (B) Calcoflour white staining for detection of seed coat phenotype variation amongst WT, *MIR775*OXP, SALK_151601.

## 5. Discussion

MicroRNAs have evolved as subtle regulators of gene expression. Recently, a few non-conserved miRNAs having a significant role in plant growth and development have been identified. The miR775 is considered to be a non-conserved, newly evolved miRNA in *A. thaliana* (Rajagopalan et al., 2006). A few studies have reported that the expression of miR775 is altered under stressed conditions (Moldovan et al., 2010; Liang et al., 2015). However, the regulatory mechanism of miR775 is still unexplored.

We observed ubiquitous expression of miR775 under standard growth condition. However, under UV and hypoxic conditions expression of miR775 was increased. The analysis of putative promoter sequence of *MIR775* indeed suggested presence of different stress responsive elements including light and/or UV light which are prevalent in high altitude areas. Recently, Zhang et al. (2021) reported that HY5 negatively regulates the expression of miR775. However, we did not observe down-regulation of miR775 under experimental light conditions. The high altitude populations are exposed to high UV light and other abiotic stress conditions (Körner, 2007). The increased expression of miR775 might be an adaptive strategy of the plants in high altitude areas. Besides light related motifs, we also observed enrichment of hypoxia related motifs in the *MIR775* promoter sequence. Thus, as expected the expression of miR775 was elevated under hypoxic condition as compared to the control. Earlier study, while analysing hypoxia responsive miRNAs, also observed increased expression of miR775 under hypoxic condition (Moldovan et al., 2009). The partial atmospheric pressure is generally less in high altitude areas as compared to sea level. It is reported that hydrogen peroxide is involved in hypoxia signalling and thereby regulating response to hypoxia (Blokhina et al., 2001; Vergara et al., 2012; Yang, 2014). Moreover, a genome wide study also indicated down-regulation of the validated target gene of miR775 (discussed below) under hypoxic condition (Branco-Price et al., 2005). Thus we observed more accumulation of hydrogen peroxide in the target mutant and *MIR775*OXP as compared to the WT. It will be interesting to investigate the regulatory mechanisms of miR775 that might aid high altitude populations to adapt under hypoxic condition.

miRNAs exerts their effect by regulating target genes. Our *in-silico* target analyses predicted a *galactosyltransferase* gene as the target of miR775 which was further confirmed experimentally using modified 5'RLM-RACE. Zhang et al. (2021) also reported the same gene as the target of miR775. However, while they reported cleavage of target as unconventional, being deviated from conventional cleavage site between 10^th^ and 11^th^ position of complementarity, our data shows perfect cleavage of the target by the miR775. Galactosyltransferases are reported to participate in the assembly of various complex polysaccharides and proteoglycans by adding galactose to various glycol-conjugates (Velasquez et al., 2011; Qin et al., 2017)). Further, galactosyltransferases are reported to be involved in growth and developmental processes (Velasquez et al., 2011; Geshi et al., 2013; Basu et al., 2015a; Basu et al., 2015b; Showalter and Basu, 2016; Qin et al., 2017; Suzuki et al., 2017). Our phylogenetic analysis suggested the target GALT is a probable β-(1,3)-galactosyltransferase. The biological role of different β-(1,3)-galactosyltransferases have been reported. For example, *GhGalT1*, a *β-(1,3)-galactosyltransferase* in cotton plays critical role in regulating cotton fibre development by controlling the glycosylation of AGPs (Qin et al., 2017) and *KNS4/UPEX1*, another GT31 family member is reported to be involved in regulating arabinogalactan biosynthesis and pollen development in *A. thaliana* (Suzuki et al., 2017). Differential staining by β-Gal-Yariv reagent, a phenylglycoside that is known to specifically bind to the β-(1,3)-galactan chains (Kitazawa et al., 2013) suggests that the target gene of miR775 is likely involved in regulating the β-(1,3)-galactan content of arabinogalactans in the stem. Besides, it is also reported that GT31 family members govern the properties of seed coat mucilage by regulating formation of complex carbohydrate (Basu et al., 2015a; Basu et al., 2015b). Differential staining of seed coat of the target mutant and WT plants with calcofluor white also suggests the target GALT might affect seed coat mucilage.

Recently, it has been reported that miR775 may regulate cell and organ size in *A. thaliana* by cell wall remodelling involving complex polysaccharides (Zhang et al., 2021). In our study, differential staining of stem cross-section and seed coat between target mutant and WT also suggest differential composition of cell wall components. Differential staining of stem cross-section and/ or seed coat between target mutant and WT also suggest differential composition of complex polysaccharides in these organs.

Although we did not carry out detailed mechanisms, but our phenotypic assessment of the target mutant and *MIR775*OXP also suggest putative role of miR775 in plant growth and development, particularly in regulating rosette size. Importantly, the effect of target down-regulation was more prominent in vegetative growth stage as compared to the reproductive traits. Moreover, as expected the knockout effect of the target gene on manifestation of traits was more prominent as compared to the down-regulation of the target by miR775. This was clearly evident when there was no difference in trait manifestation between *MIR775*OXP and the WT. For example, there was no difference in silique length and seed number per silique or β-Gal-Yariv staining intensity in stem cross-section between *MIR775*OXP and WT, although there were clear differences between the knockout mutant and WT. It may be that the relative stoichiometry of miR775/target is not quantitatively enough for such trait manifestation.

## 6. Conclusions

We describe the functional significance of miR775, an evolutionary young miRNA that targets a probable *β-(1,3)-galactosyltransferase*. Both, miR775 and its target gene are ubiquitously expressed having a more pronounced role during vegetative growth leading to increased rosette area. Increased expression of miR775 under UV light and hypoxia intuitively suggest it might aid in high altitude adaptation of plants, since these are more prevalent in these area. However, detailed studies need to be conducted to decipher any role of miR775 in plant adaptation in high altitude areas. Nevertheless, our findings high lights the significance of a young miRNA in plant growth and development. Since miR775 is only reported in *A. thaliana* while galactosyltransferases are conserved, the significance of newly evolved young miRNAs in regulating galactosyltransferases needs to be analysed in other plant species.

## Acknowledgment

The authors acknowledge the financial support of Council of Scientific and Industrial Research (CSIR), New Delhi and partly by Department of Biotechnology (DBT), Government of India, New Delhi, India. PM and AKV also acknowledge the University Grant Commission (UGC), New Delhi for providing the fellowship.

The institutional ethic committee reference number for the manuscript is CSIR-NBRI_MS/2021/01/05.

## Author contribution

PM, AS, AKV and RS conducted experiments. PM analysed the results and wrote the manuscript with SR. SR designed and arranged fund for the experiments.

## Supplementary Figures

**Supplementary Fig. 1**: Relative expression of *DCL1* (AT1G01040) in *MIR775*OXP lines and WT as determined by qRT-PCR. Error bars represent ±SE with n=3. ns indicates non-significant variation in OXP lines as compared to WT.

**Supplementary Fig. 2:** Relative expression of *galactosyltransferase* (AT1G53290) target gene in different tissues of WT Col-0 ecotype as determined by qRT-PCR. Bars represent ±SE (n=3). *** indicate values which were significantly different with P value <0.001 (Student’s t-test).

**Supplementary Fig. 3:** qRT-PCR for relative expression of AT1G53290 in SALK_151601 line as compared to WT. Error bars represent ±SE (n=3). *** indicates significant difference from WT with p value < 0.001 (Student’s t-test).

**Supplementary Fig. 4:** Response of WT, *MIR775*OXP and target mutant (SALK_151601) in terms of hydrogen peroxide accumulation after six hours of treatment with mitochondrial inhibitors, as determined by (A) DAB (3,3′-Diaminobenzidine) staining (B) hydrogen peroxide content estimation in microgram per gram fresh weight (μg/gFW). Errors bars indicate ± SE (n=3), while *, ** indicates significant difference between treatment and control with p value < 0.05, <0.01 (Student’s t-test).

**Supplementary Fig. 5:** Protein association network of AT1G53290 adapted from STRING (Search Tool for the Retrieval of Interacting Genes/Proteins) (https://string-db.org/). The target gene is predicted to associate with many other proteins. Nodes in the network represent different proteins with the target protein of miR775 in the center, while edges represent the protein-protein association. Yellow edges represent association curated on the basis on text mining, while pink represent experimentally determined associations.

**Supplementary Fig. 6:** Staining with β-Gal Yariv reagent in (A) stem sections of WT and *MIR775*OXP (B) in pollen of WT, *MIR775*OXP and target mutant (SALK_151601).

## Supplementary Tables

**Supplementary Table 1:** List of primer sequences used

**Supplementary Table 2:** Putative targets of miR775 as predicted by psRNATarget prediction tool

**Supplementary Table 3:** List of motifs found in the promoter sequence of *MIR775*

**Supplementary Table 4:** Results of statistical analysis for morphological traits

**Supplementary Table 5:** Details of proteins predicted to associate with target GALT as identified by STRING

