## Supplementary Figures for "MicroRNA775 targets a *β-(1,3)-galactosyltransferase* to regulate growth and development in *Arabidopsis thaliana*"

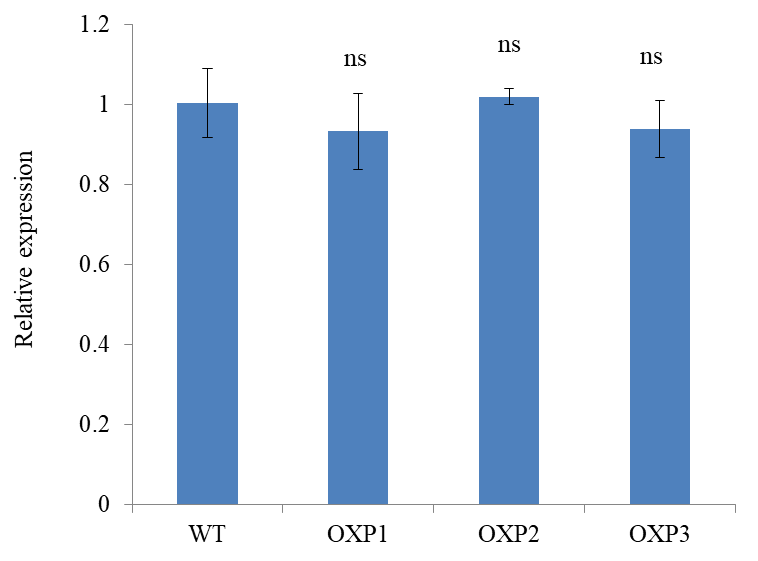

**Supplementary Fig. 1:** Relative expression of *DCL1* (AT1G01040) in *MIR775*OXP lines and WT as determined by qRT-PCR. Error bars represent ±SE with n=2. ns indicates non-significant variation in OXP lines as compared to WT.

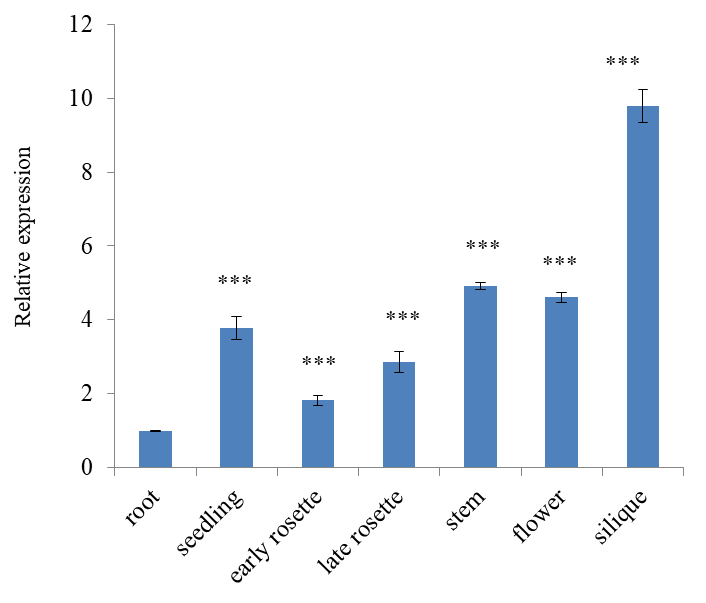

**Supplementary Fig. 2:** Relative expression of *galactosyltransferase* (AT1G53290) target gene in different tissues of WT Col-0 ecotype as determined by qRT-PCR. Bars represent ±SE (n=3). *** indicate values which were significantly different with P value <0.001 (Student’s t-test).

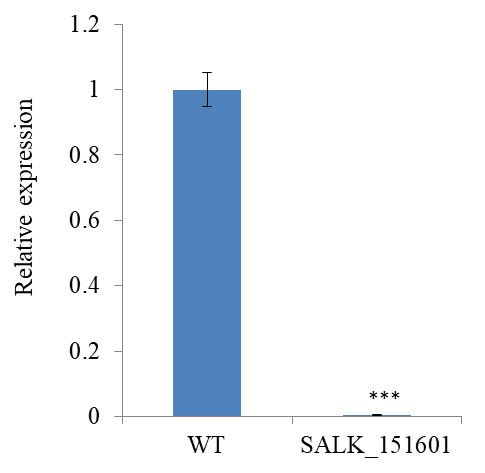

**Supplementary. Fig. 3:** qRT-PCR for relative expression of AT1G53290 in SALK_151601 line as compared to WT. Error bars represent ±SE (n=3). *** indicates significant difference from WT with p value < 0.001 (Student’s t-test).

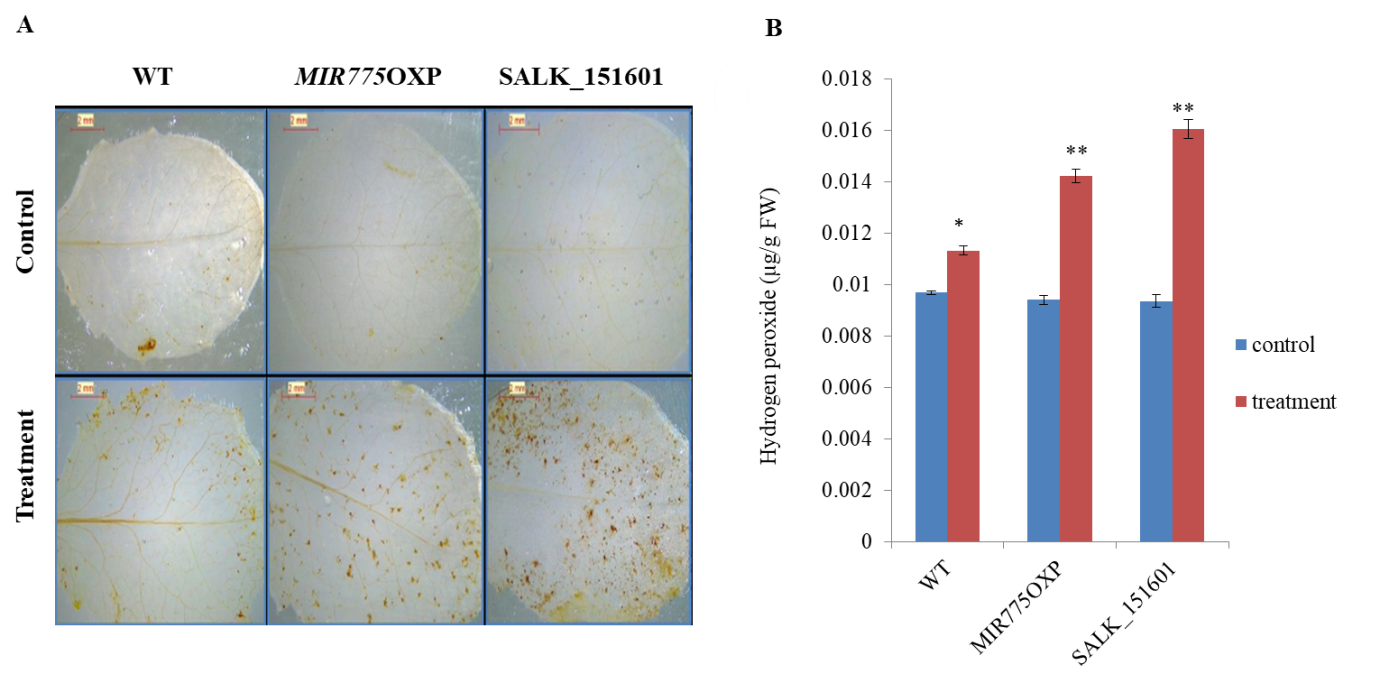

**Supplementary Fig. 4:** Response of WT, *MIR775*OXP and target mutant (SALK_151601) in terms of hydrogen peroxide accumulation after six hours of treatment with mitochondrial inhibitors, as determined by (A) DAB (3,3′-Diaminobenzidine) staining (B) hydrogen peroxide content estimation in microgram per gram fresh weight (µg/gFW). Errors bars indicate ± SE (n=3), while *, ** indicates significant difference between treatment and control with p value < 0.05, <0.01 respectively (Student’s t-test).

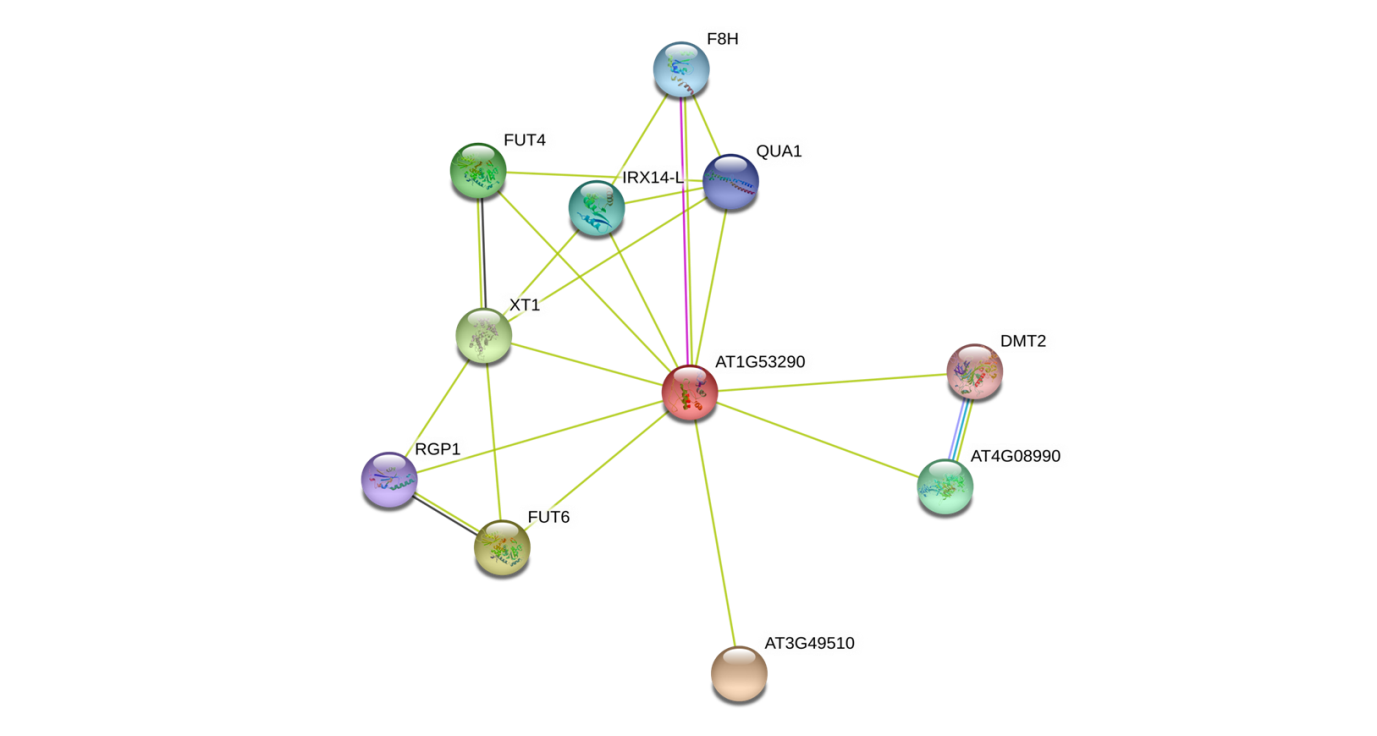
**Supplementary Fig. 5:** Protein association network of AT1G53290 adapted from STRING (Search Tool for the Retrieval of Interacting Genes/Proteins) (https://string-db.org/). The target gene is predicted to associate with many other proteins. Nodes in the network represent different proteins with the target protein of miR775 in the center, while edges represent the protein-protein association. Yellow edges represent association curated on the basis on text mining, while pink represent experimentally determined associations.

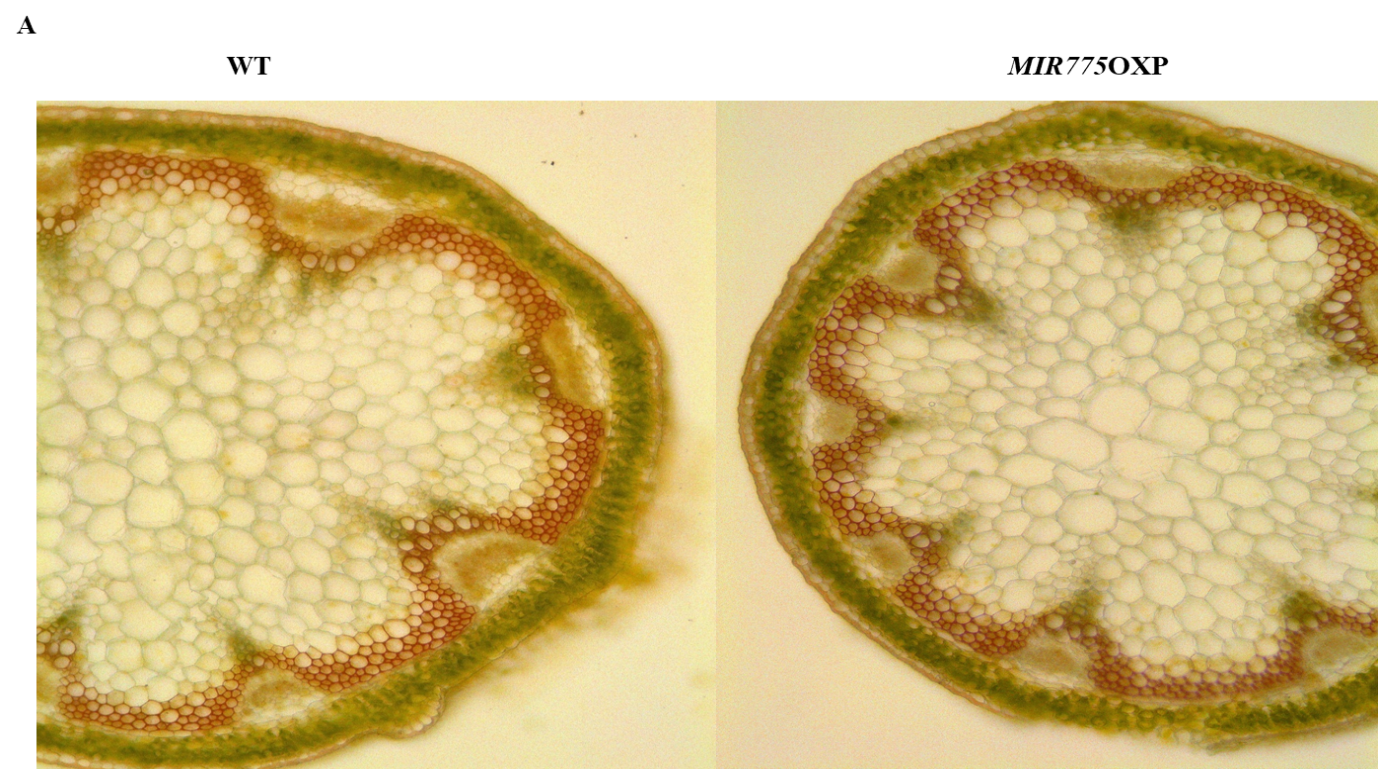

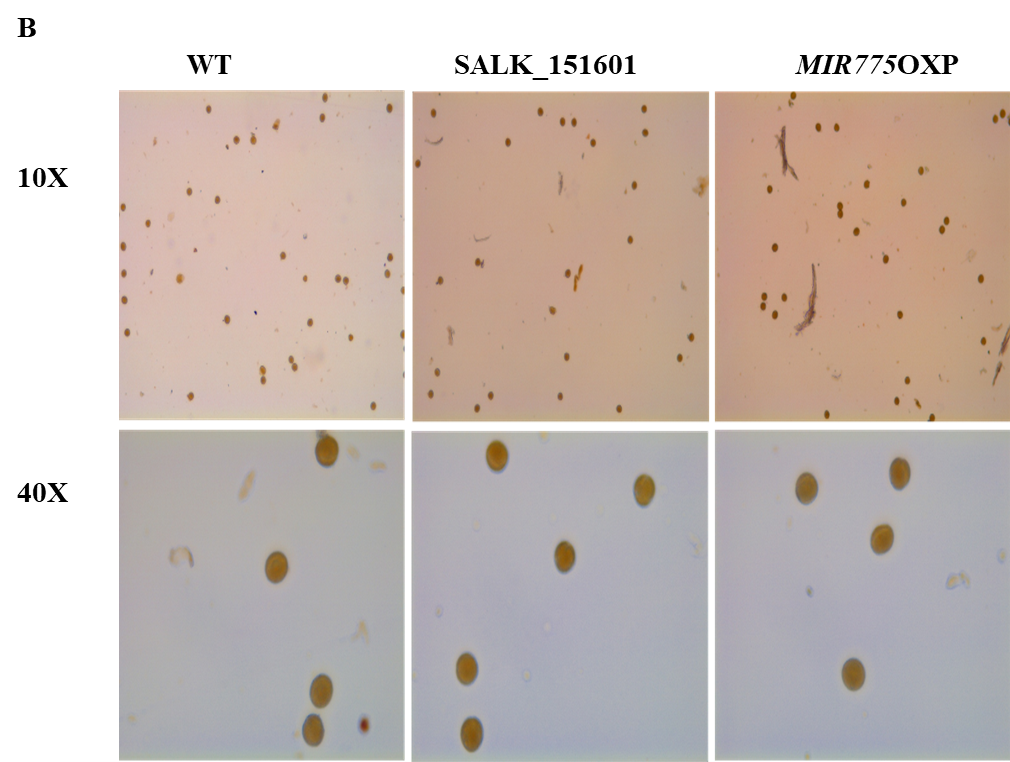

**Supplementary Fig. 6:** Staining with β-Gal Yariv reagent in (A) stem sections of WT and *MIR775*OXP (B) in pollen of WT, *MIR775*OXP and target mutant (SALK_151601).
